# PhyB induces intron retention and uORF-mediated translational inhibition of PIF3

**DOI:** 10.1101/813840

**Authors:** Jie Dong, Haodong Chen, Xing Wang Deng, Vivian F. Irish, Ning Wei

**Affiliations:** Department of Molecular, Cellular and Developmental Biology, Yale University, New Haven, Connecticut 06520, USA; State Key Laboratory of Protein and Plant Gene Research, Peking-Tsinghua Center for Life Sciences, School of Advanced Agricultural Sciences and School of Life Sciences, Peking University, Beijing 100871, China

**Keywords:** phytochrome B, PIF3, translational regulation, alternative splicing, intron retention, uORF, photomorphogenesis, short day, diurnal cycle, light regulation, Arabidopsis

## Abstract

The phytochrome B (phyB) photoreceptor stimulates light responses in plants in part by inactivating repressors of light responses such as phytochrome-interacting factor 3 (PIF3). It has been established that activated phyB inhibits PIF3 by rapid protein degradation and decreased transcription. PIF3 protein degradation has been shown to be mediated by EIN3-BINDING F-BOX PROTEIN (EBF) and LIGHT-RESPONSE BTB (LRB) E3 ligases, the latter simultaneously targeting phyB for degradation. In this study, we show that PIF3 level is additionally regulated by alternative splicing and protein translation. Overaccumulation of photo-activated phyB, which occur in the mutant defective for *LRB* genes under continuous red light (Rc), induces a specific alternative splicing of *PIF3* that results in retention of an intron in the 5’UTR of *PIF3* mRNA. In turn, the upstream opening reading frames (uORF) contained within this intron inhibit PIF3 protein synthesis. The phyB-dependent alternative splicing of *PIF3* is diurnally regulated under the short-day light cycle. We hypothesize that this reversible regulatory mechanism may be utilized to fine-tune the level of PIF3 protein in light-grown plants, and may contribute to the oscillation of PIF3 protein abundance under the short-day environment.

**One Sentence Summary:** Light down-regulates PIF3 by multiple mechanisms. We show that phyB induces an alternative splicing event that inhibits PIF3 protein translation, and that is regulated by short-day diurnal cycle.

## Introduction

The phytochromes (phy) are important photoreceptors regulating plant growth and development throughout its lifecycle. Among the five members of the phytochrome family in *Arabidopsis*, phyB is relatively abundant in light and plays a major role in controlling hypocotyl growth under prolonged continuous red light (Rc) (Li et al., 2011). Phytochrome-Interacting Factors (PIFs) are basic helix-loop-helix transcription factors that function to repress light responses (Leivar et al., 2008; Shin et al., 2009; Pham et al 2018). PIF3, the first characterized member of the PIF family (Ni et al., 1998), accumulates abundantly in dark-grown seedlings to maintain etiolation, and it rapidly declines upon light exposure, allowing establishment of photomorphogenesis in seedlings (Bauer et al., 2004; Zhang et al., 2013). During de-etiolation, photoactivated phyB directly binds to PIF3 and induces PIF3 phosphorylation, which are necessary for the 26S proteasome-mediated degradation of PIF3 (Al-Sady et al., 2006; Ni et al., 2013). In addition, phyB down-regulates *PIF3* transcription by about 3-fold during de-etiolation and interferes with PIF3 DNA binding ability (Shi et al., 2016; Park et al., 2018). Inhibition of PIF3 at the level of translation has not been reported, although light signals generally promote translation activity globally (Paik et al., 2012; Chen et al., 2018).

After seedling establishment, normal light-grown plants maintain steady-state lower expression of PIFs, which continue to modulate plant development in response to light and temperature (Leivar and Monte, 2014; Pham et al., 2018). For examples, PIF3 was shown to play prominent roles in promoting dark-period elongation growth under short-day conditions (Soy et al., 2012; 2016), and in freezing tolerance (Jiang et al., 2017).

Studies on phyB induced rapid protein degradation of PIF3 have revealed that PIF3 can be degraded through at least two pathways. During de-etiolation, F-box proteins EIN3-BINDING F BOX PROTEIN (EBF) mediate PIF3 degradation via SCF^EBFs^ ubiquitin E3 ligases to facilitate photomorphogenesis of plants (Dong et al., 2017). Plants deficient for EBFs exhibit reduced light sensitivity and inefficient de-etiolation. Under high fluence red light, PIF3 can also be degraded via LIGHT-RESPONSE BTBs (LRBs), the ubiquitin E3 ligases (CRL3^LRBs^) that target phyB for degradation to attenuate light responses (Ni et al., 2014). Plants deficient for LRBs show hypersensitivity to light in a phyB dependent manner (Christians et al., 2012; Ni et al., 2014; Dong et al., 2017).

To study the functional dynamics between EBFs and LRBs in the regulation of PIF3, we eliminated both types of PIF3 E3 ligases and generated a *ein3 ebf lrb* mutant line. Using this mutant line, we uncovered an unexpected mechanism by which phyB inactivates PIF3: i.e. inhibiting PIF3 protein translation via alternative splicing (AS).

## Results and Discussion

To knock-out both ubiquitin E3 families of PIF3, we generated *ein3 ebf lrb* hextuple mutant by crossing the viable *ein3 ebf1 ebf2* (*ein3 ebf*) mutant with *lrb1 lrb2 lrb3* (*lrb*) (Figure 1). These mutants were further confirmed to lack the corresponding gene transcripts in RT-PCR (Supplemental Figure 1). The resulting phenotypes under continuous red light (Rc) showed that the *ein3 ebf* mutant exhibited longer hypocotyls compared to *ein3*, while *lrb* showed shorter hypocotyls compared to Col (Figure 1A and 1B), consistent with previous observations (Dong et al., 2017; Ni et al., 2014; Christians et al., 2012). The *ein3 ebf lrb* mutant almost phenocopied *lrb* in having a shortened hypocotyl under Rc, suggesting that *lrb* strongly suppressed *ein3 ebf* (Figure 1A and 1B). The hyper-photosensitive phenotype of *ein3 ebf lrb* seemed at odds with the anticipated accumulation of PIF3 in high levels, and prompted us to examine the PIF3 protein levels in these mutants.

**Figure 1.**
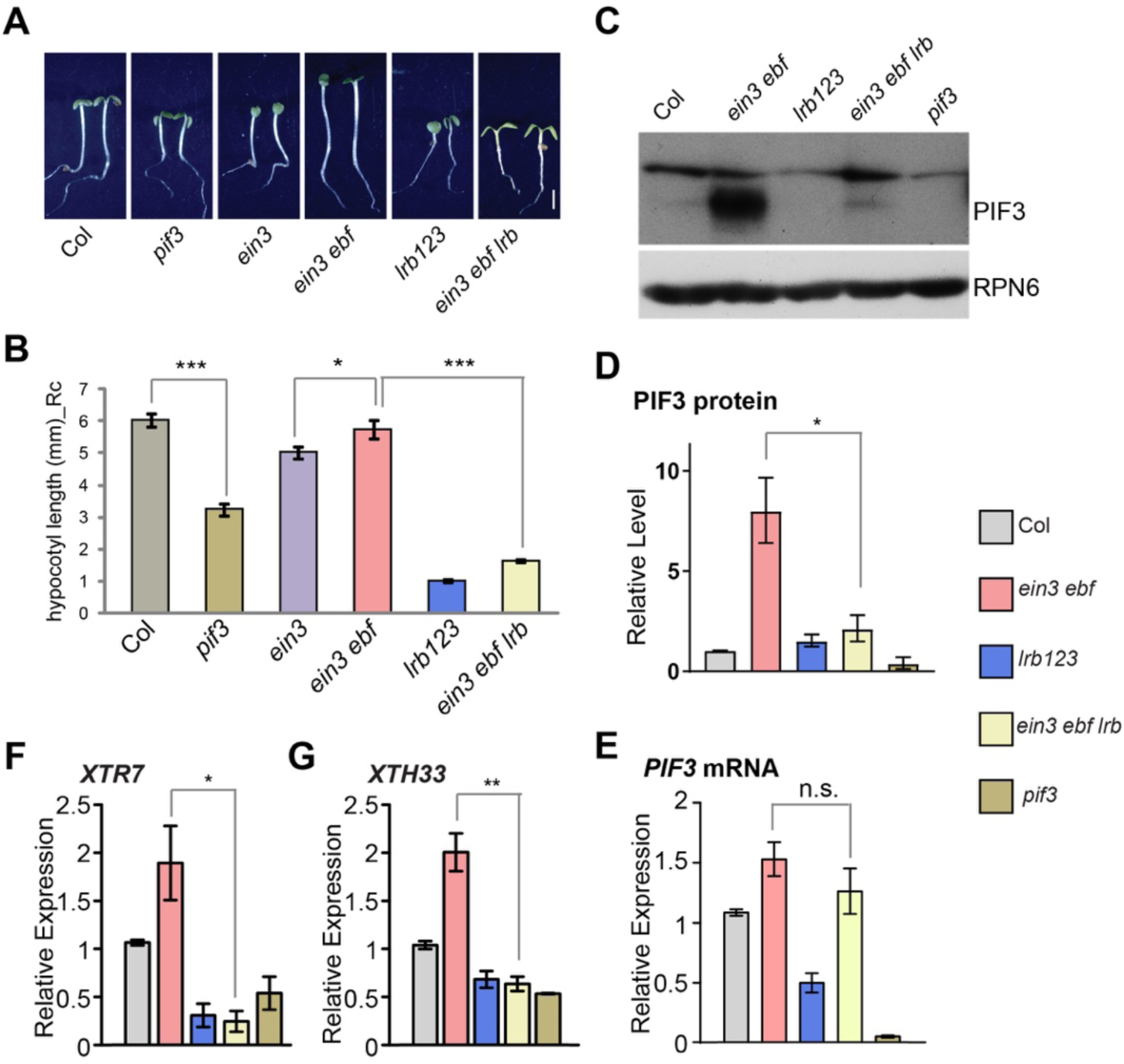
*LRB* mutations resulted in a decrease of PIF3 protein and hypocotyl elongation in *ebf* mutant background under Rc. (**A**) Representative images of Col, *pif3*, *ein3*, *ein3 ebf1 ebf2* (*ein3 ebf*), *lrb1 lrb2 lrb3* (*lrb123*) and *ein3 ebf1 ebf2 lrb1 lrb2 lrb3* (*ein3 ebf lrb*) seedlings grown under 10 μmol/m^2^/s continuous red light (Rc) for 4 days. Scale bar= 2mm. EIN3 is an ethylene response factor and a degradation target of EBF (Shi et al., 2016). Note: to offset the ethylene phenotype, the *ebf* mutant is represented by *ein3 ebf* with *ein3* as its control. (**B**) Mean hypocotyl lengths of each genotype shown in (**A**). Data were shown in mean±SEM. (**C and D**) Accumulation of PIF3 protein in *ein3 ebf* was suppressed in *ein3 ebf lrb*. Proteins were extracted from 4-day-old seedlings of indicated genotype grown under Rc, and western blots were performed using anti-PIF3 and anti-RPN6 antibodies. A representative western blot was shown in (**C**), and the mean relative PIF3 protein level in each genotype was shown in (**D**). Data were shown in mean±SEM of 3 biological replicates. Transcript levels of *PIF3* (**E**), *XTR7* **(F)**, and *XTH33* **(G)** in each genotype under Rc. Total RNAs were extracted from 4-day-old seedlings grown under Rc and reverse transcribed for RT-qPCR analyses. *ACT2* was used as internal control. Data were shown in mean±SD of 3 biological replicates. All statistic significances were calculated by Student’s *t* test: n.s., p>0.05; *, p<0.05; ***, p<0.001.

By eliminating both E3 ubiquitin ligase families that regulate PIF3 abundance, one would expect that PIF3 protein would be stabilized in the *ein3 ebf lrb* mutant. However, western blot showed that PIF3 protein accumulated only in *ein3 ebf*, but not in *lrb* nor in *ein3 ebf lrb* mutants (Figure 1C and 1D). PIF3 protein levels were much higher in *ein3 ebf* than in *ein3 ebf lrb*, while *PIF3* transcript levels were comparable between these two mutant lines (Figure 1D and 1E). In addition, PIF3 activities matched with PIF3 protein levels in these mutants, based on the expression of the PIF3 target genes *XTR7* and *XTH33* (Figure 1F and 1G). In the dark, relative levels of PIF3 protein corresponded with their respective mRNA levels among the mutants tested (Supplemental Figure 2), suggesting the regulation of PIF3 protein levels by LRBs and EBFs are light-dependent. Taken together, the *lrb* mutations appeared to have suppressed the PIF3 over-accumulation of *ein3 ebf* by decreasing PIF3 protein expression without affecting its transcript levels, and one possibility is that *lrb* could potentially inhibit translation of PIF3 under Rc. As *lrb* mutations primarily cause phyB accumulation and light hypersensitivity (Christians et al., 2012; Ni et al., 2014), we hypothesized that phyB may negatively regulate PIF3 translation.

Light causes wide-spread alternative splicing (AS) in plant transcriptome (Shikata et al., 2014; Hartmann et al., 2016; Xin et al., 2017; Godoy Herz et al., 2019). Although *PIF3* has not been experimentally demonstrated to undergo light regulated AS, based on the information from Araport (Arabidopsis Information Portal: www.araport.org), transcripts with alternative splicing patterns at an intron of approximately 500 bp in *PIF3* 5’ untranslated region (UTR) have been found in the database (Figure 2A). We refer to this intron as Intron_AS_ in this study, and monitored its splicing level by RT-qPCR. We found that the Intron_AS_-retained form was significantly increased relative to total *PIF3* mRNA in Col under Rc, as compared to dark (Dc; Figure 2B), suggesting that Intron_AS_ retention in *PIF3* mRNA is a light-dependent event. Furthermore, mutation of phyB essentially abolished the light (Rc) induced increase of Intron_AS_ retention (Figure 2B), indicating that phyB is critical in red light responsive Intron_AS_ retention in *PIF3* mRNA. Moreover, *lrb* mutations caused higher Intron_AS_ retention under Rc (Supplemental Figure 3), which is consistent with higher levels of phyB accumulation in *lrb* (Christians et al., 2012; Ni et al., 2014).

**Figure 2.**
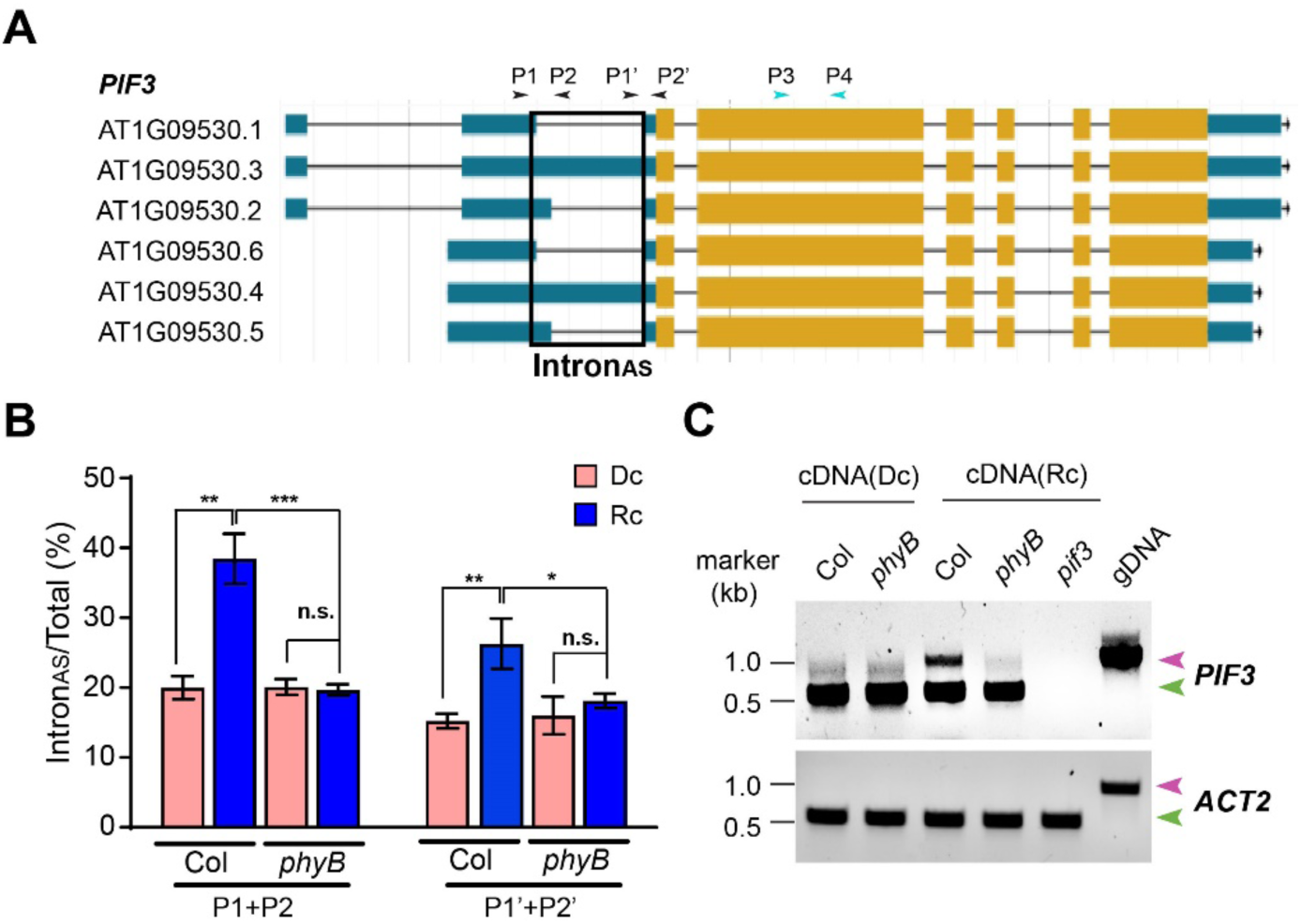
Light induces Intron_AS_ retention in *PIF3* mRNA via phyB. (**A**) The schematic diagram of various *PIF3* pre-mRNAs (Araport) and the primer pairs used for RT-qPCR. (**B**) The relative percentages of Intron_AS_ retained *PIF3* mRNA (represented by P1+P2 or P1’+P2’) to total *PIF3* mRNA (represented by P3+P4) in Col and *phyB-9* (*phyB*). cDNA was made from seedlings grown in the dark (Dc) or in 10 µmol/m^2^/s continuous red light (Rc) for 4 days. The RT-qPCR data are shown as the mean±SD of 3 biological replicates. Statistic significances were calculated by Student’s *t* test: n.s., p>0.05; *, p<0.05; **, p<0.01; ***, p<0.001. (**C**) The visualization of Intron_AS_ retention in PIF3 mRNA by PCR gel imaging. Primers used for *PIF3* are P1 and P4 shown in (**A**). Primers used for *ACT2* are ACT2I1gelf and ACT2I1gelr. Arrows in pink and green represent PCR products with and without Intron_AS_ retention, respectively.

To better visualize the retention of Intron_AS_, we performed gel electrophoresis of the PCR products using a primer pair that would produce a longer product when Intron_AS_ is retained, and a shorter product when Intron_AS_ is spliced (Figure 2C). The *ACTIN2* (*ACT2*) primer pair produced only the short-sized band from all cDNA samples while produced a long-sized band from genomic DNA (Figure 2C), suggesting there was no intron retention in *ACT2* mRNA in those samples. By contrast, the *PIF3* primer sets produced both the intron-containing and intron-spliced bands from the cDNA samples. The Intron_AS_-retained form was strongly enriched under Rc compared to Dc in Col, but the light-induced intron retention was nearly absent in *phyB-9*. This result shows again that Intron_AS_ retention in *PIF3* mRNA occurred under red light, and that this event was dependent on phyB (Figure 2C). As an additional control, we examined other introns in *PIF3* mRNA and found no evidence of retention of any of other introns (Supplemental Figure 4). These data show that photoactivated phyB can induce an alternative splicing event that results in the retention of Intron_AS_ in *PIF3* mRNA.

Extensive studies have shown that the 5’UTR can influence its mRNA translation (Araujo et al., 2012; von Arnim et al., 2014). Since Intron_AS_ resides within the 5’UTR of *PIF3* mRNA, we next asked whether the retention of Intron_AS_ may affect PIF3 translation. We generated T7 promoter-driven constructs in which either normally spliced *PIF3* 5’ UTR (5U) or Intron_AS_-retained 5’UTR (5IU) was inserted upstream of the *PIF3* coding sequence. In a mammalian cell-based *in vitro* translation assay using either DNA or RNA as template, 5U produced higher levels of PIF3 proteins than did 5IU (Figure 3A). Clearly, retaining the Intron_AS_ sequence in *PIF3* 5’UTR inhibited protein translation even in a heterologous *in vitro* translation system.

**Figure 3.**
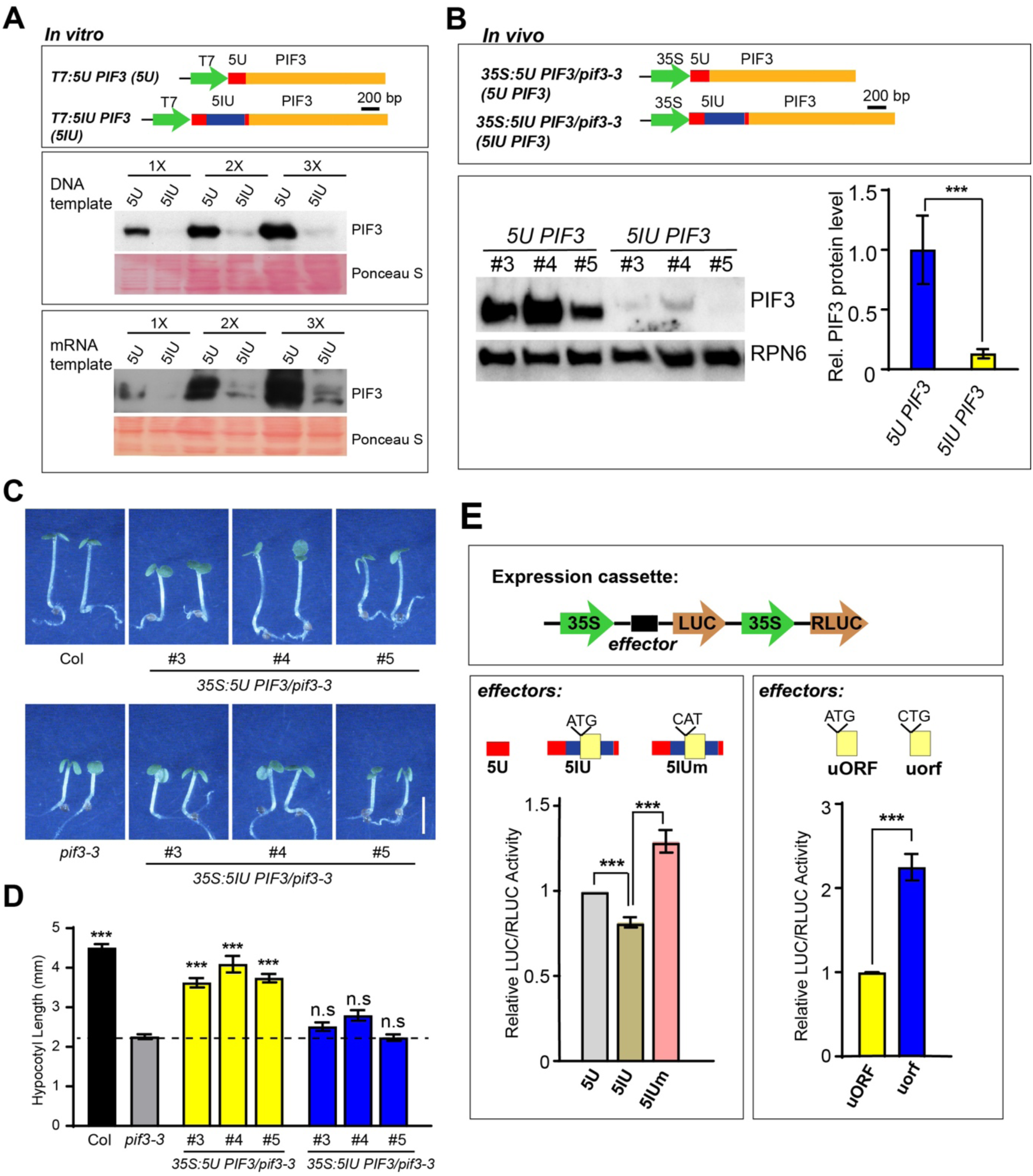
Intron_AS_ retention inhibits *PIF3* mRNA translation via uORFs. (**A**) Intron_AS_ retention in *PIF3* 5’UTR inhibits protein translation *in vitro*. The schematic diagram shows the templates used for *in vitro* translation. DNA or *in vitro* transcribed mRNA was used as template for in vitro translation, and products were analyzed by anti-PIF3 western blots. 1×, 2× and 3× indicates the fold increasing amounts of *in vitro* translation samples that were loaded. (**B**) Intron_AS_ in the *PIF3* 5’UTR inhibits PIF3 protein production *in vivo*. The schematic diagram shows the expression cassettes used for generating transgenic plants in the *pif3-3* background. Proteins extracted from 3-day dark grown seedlings were analyzed by anti-PIF3 western blots, with RPN6 as a loading control. Mean relative PIF3 protein levels were normalized to the corresponding RPN6 control and total *PIF3* mRNA level. Data are shown as the mean±SD of 3 independent transgenic lines. The value in *5U PIF3* transgenic plants was set as 1.0. (**C** and **D**) *5U PIF3* transgene, but not *5IU PIF3*, could rescue the phenotypes of *pif3-3* under Rc. Three-day-old seedlings grown under Rc were photographed (**C**), and hypocotyl length was measured (**D**). Data were shown as the mean ± SEM of at least 30 seedlings. Statistical significance was calculated against *pif3-3* using Student’s *t* test: n.s., p>0.05; ***, p<0.001. (**E**) The constructs carrying indicated effectors were expressed in Arabidopsis protoplasts, and the ratio of LUC activity to RLUC activity was determined. Data were shown as the mean±SD of 4 biological replicates. All statistical significances were calculated by Student’s *t* test: ***, p<0.001.

To test the effect of Intron_AS_ *in planta*, we generated transgenic plants in a *pif3-3* mutant background that carried the following transcription cassettes under the 35S promoter: *5U PIF3* or *5IU PIF3*, in which normally spliced 5’UTR or Intron_AS_-containing 5’UTR, respectively, were placed upstream of PIF3 coding region (Figure 3B). Three independent *5U PIF3* and *5IU PIF3* transgenic lines with similar levels of total *PIF3* mRNA transcript were selected, and Intron_AS_-containing *PIF3* transcript was confirmed to exist only in *5IU PIF3* transgenic lines (Supplemental Figure 5). It appeared that, in the context of this artificial Intron_AS_-containing *PIF3* transgene, Intron_AS_ could be spliced but very inefficiently, as most of the transcript retained the intron (Supplemental Figure 5C). In darkness when PIF3 protein was stable, western blot showed that PIF3 protein expressed in *5U PIF3* lines, but was almost undetectable in *5IU PIF3* lines (Figure 3B). Quantification showed that retention of Intron_AS_ in *PIF3* 5’UTR caused an approximately 80% reduction in PIF3 protein levels (Figure 3B). These results indicate that forced retention of Intron_AS_ upstream of the PIF3 coding sequence inhibits the protein production. As a result, the *5IU PIF3* transgene was unable to restore the phenotype of *pif3-3* in hypocotyl elongation under Rc, in contrast to the rescue seen with *5U PIF3* (Figure 3C and 3D). We deduce from these results that the retention of Intron_AS_ in the 5’UTR of *PIF3* mRNA can inhibit PIF3 protein expression both *in vitro* and *in vivo*, consequently affecting PIF3-regulated hypocotyl elongation.

Upstream Open Reading Frame (uORF)-mediated translation inhibition has been shown to regulate gene expression involved in light response as well as other developmental and metabolic processes in plants (von Arnim et al., 2014; Kurihara et al., 2018). The *PIF3* Intron_AS_ sequence contains multiple AUG start codons and putative uORFs (Supplemental Figure 6). The longest uORF, which is 129 nucleotides in length just upstream of the PIF3 main ORF, is considered the most likely candidate uORF to inhibit downstream PIF3 translation, even though the start codon context is suboptimal (Kozak, 2001). To test this idea, we carried out a dual-luciferase protoplast transient assay where the 5’UTR testing effector was inserted upstream of firefly luciferase (LUC) reporter, with a 35S promoter driven Renilla luciferase (RLUC) as the internal control (Figure 3E). The testing effectors include: *PIF3* 5’UTR (5U), Intron_AS_-containing 5’UTR (5IU), or Intron_AS_ with the mutated form of uORF start codon 5I_AS_U (5IUm). In another set, we explicitly tested the effect of the uORF without other Intron_AS_ sequences: either the uORF with an ATG start codon, or mutated uORF with CTG (uorf). The data showed that relative LUC activity in 5IU and uORF was significantly lower than that in 5IUm and uorf, respectively (Figure 3E). These results indicate that the 129 nt uORF of Intron_AS_ can inhibit the translation of the downstream ORF.

We noticed that the inhibitory effect of Intron_AS_-inclusion was not as strong in this transient assay (Fig. 3E 5U vs 5IU) compared to the *in planta* assay (Fig. 3B), which was probably due to the difference in the assay system. In addition, the dramatic inhibition of protein expression seen in the experiments of Figure 3A and 3B could be caused by the combination of the uORFs in Intron_AS_, as oppose to Figure 3E that tested the 129 nt uORF exclusively. Regardless, our data support the conclusion that retention of Intron_AS_, which contains uORFs, causes a blockage in PIF3 expression at the level of translation.

It is well established that phyB activation leads to a sharp decline in PIF3 abundance during de-etiolation (Bauer et al., 2004; Al-Sady et al., 2006; Dong et al., 2017). phyB stimulated PIF3 protein degradation accounts for predominant portion of PIF3 downregulation (Al-Sady et al., 2006), while decreased transcription of *PIF3* also occurs (Shi et al., 2016). Here, our data show that phyB can additionally inhibit PIF3 protein translation via retention of Intron_AS_ under prolonged red light. However, questions remain as to how important the translation control is relative to the control of protein stability and transcription, and under what natural circumstances would the plants utilize this type of regulatory mechanism. To this end, we examined the kinetics of Intron_AS_ retention rate during de-etiolation. When dark-grown seedlings were irradiated with red light, PIF3 protein level rapidly and sharply declined within minutes and dropped more than 10 fold within hours after light exposure, while the increase of Intron_AS_ retention was more evident in prolonged light-grown plants than during the 24 hours of de-etiolation (Supplemental Figure 7). This result suggests that phyB-induced AS is unlikely to play a major role in de-etiolation, but maybe a feature associated with light-grown plants. As most light-dependent AS profiling studies were performed under the de-etiolation condition (Shikata et al., 2014; Hartmann etal., 2016; Kurihara et al., 2018), this maybe a reason why PIF3 has not been reported to undergo light-induced AS.

In growth plants, PIF3 protein abundance has been shown to oscillate under short day diurnal cycle to promote pre-dawn hypocotyl growth (Soy et al., 2012; 2016). Interestingly, unlike PIF4 and PIF5 whose transcription is strongly diurnally regulated, transcription regulation of *PIF3* is minimum under this condition, while PIF3 protein accumulates in the dark period and decline during the day (Soy et al., 2012). We examined *PIF3* Intron_AS_ retention in plants grown in short-day light cycle. The data showed that the Intron_AS_ retention rate of *PIF3* oscillated diurnally, rising post-dawn and dropping around dusk (Figure 4A). This pattern would correspond with a reduced PIF3 translation during the day, and normal protein synthesis during the night, which is consistent with peak accumulation of PIF3 protein at pre-dawn (Soy et al., 2012). We thus suggest that regulation of PIF3 protein synthesis, in coordination with PIF3 degradation, contributes to diurnal oscillation of PIF3 protein levels under short-day conditions.

**Figure 4.**
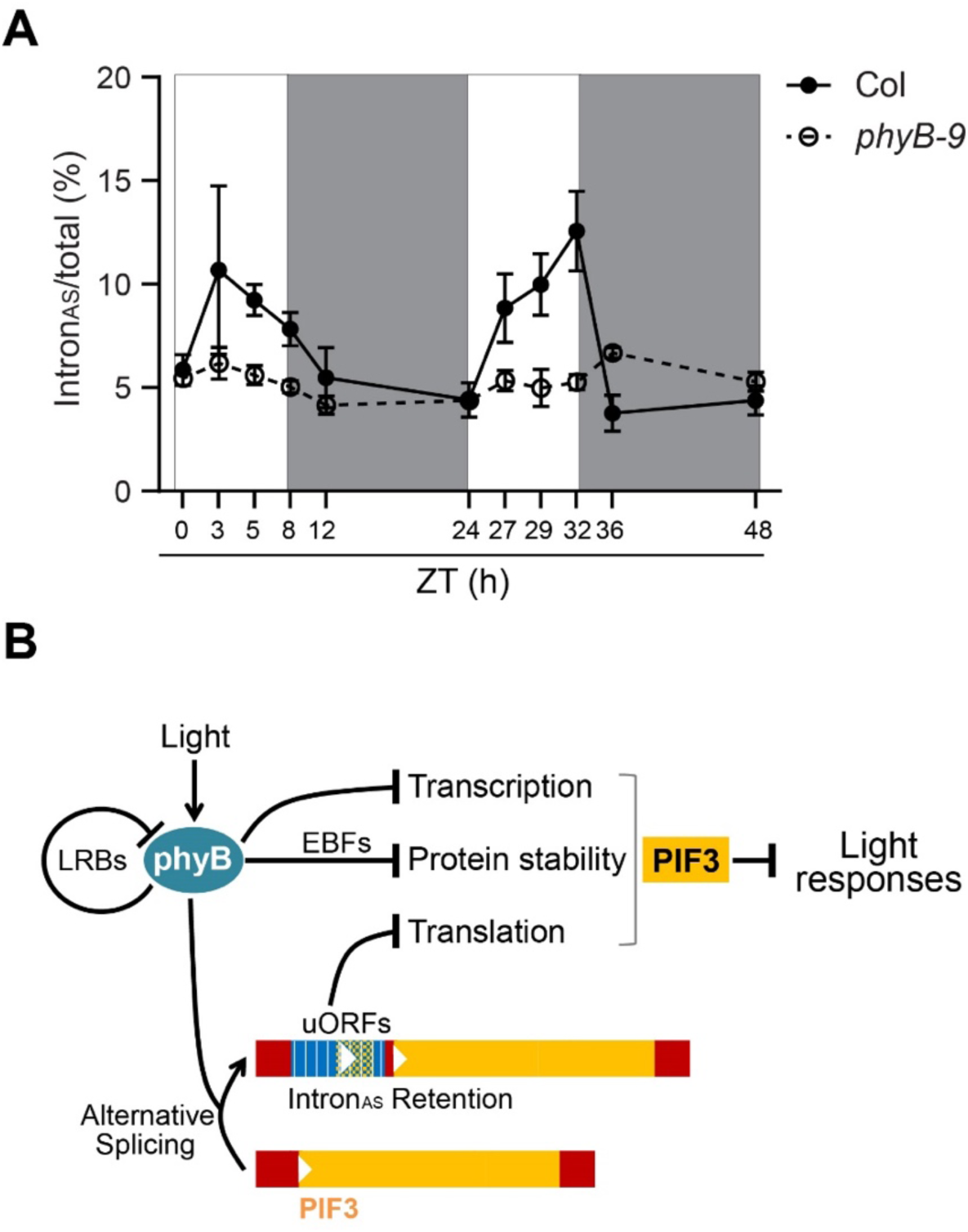
The phyB-dependent *PIF3* AS-uORF plays a role in regulating PIF3 levels, in conjunction with other regulatory mechanisms. (**A**) Diurnal regulation of Intron_AS_ retention in *PIF3* mRNA. Col seedlings were grown under short day (8-hr light/16-hr dark) condition, and samples were harvested at indicated Zeitgeber Time (ZT) from 4 day old plants. Total RNA was extracted and reverse transcribed, and cDNA was used for qPCR analysis. Data were shown in mean±sem (n=3 biological replicates). (**B**) A summary diagram showing that light-activated phyB reduces PIF3 protein level in multiple ways to stimulate light responses. In addition to triggering transcriptional down-regulation and EBF-mediated protein degradation of PIF3, phyB can also induce alternative splicing in light-grown plants that results in the retention of Intron_AS_ (shown in blue) in the 5’UTR of *PIF3* mRNA. The uORFs inside the Intron_AS_ sequence inhibit translation of PIF3 main ORF (shown in yellow). The 5’ and 3’ UTR region of *PIF3* mRNA are indicated in red. LRB E3 ligases may work as a negative feedback system to control phyB protein levels and attenuate excessive light activation.

In summary, we have identified a new regulatory mechanism of phyB on *PIF3*, and revealed that light signals can inactivate PIF3 at level of translation, apart from regulating its transcription, DNA binding, and protein stability (Figure 4B). In particular, our findings explain why the *lrb* E3 mutations can suppress the *ebf* E3 mutants phenotypically and at the level of PIF3 protein accumulation. This is because *lrb* induced phyB accumulation leads to a block of PIF3 protein synthesis, which would alleviate the problems caused by the deficient PIF3 protein degradation in *ebf* (Figure 1). It has been shown that light signals can cause the uORF inclusion in the 5’UTR of several genes as a result of global change of transcription start sites (Kurihara et al., 2018). In *PIF3*, alternative transcription initiation may also occur (Figure 2A), but it is not related to the inclusion of the uORFs. Regarding the extent of translational regulation relative to transcription and protein degradation of PIF3, we suggest that, during de-etiolation, the massive irreversible PIF3 protein degradation probably accounts for most of the rapid and dramatic drop of PIF3 levels, while the AS coupled-translational regulation is minor if it occurs at all. In grown plants, PIF3 expresses at a reduced level, and a reversible control mechanism as revealed in this study may operate to fine-tune PIF3 levels, likely in conjunction with other PIF3 regulatory mechanisms. Our data that retention of Intron_AS_ oscillates in short-day diurnal cycle support this idea. It is possible that Intron_AS_-coupled translational regulation of PIF3 may play important roles in other light modulated physiological processes in plants.

## Materials and Methods

### Plant materials and growth condition

*Arabidopsis thaliana* mutants are in the Col (Columbia) ecotype and were grown under long day conditions (16h day/ 8h dark) at 22 °C. The parental lines of the higher order mutant, *ein3 ebf1 ebf2* (*ein3 ebf*) and *lrb1 lrb2 lrb3* (*lrb*), were previously described in An et al., 2010 and Ni et al., 2014, respectively. The *pif3* mutant was described in Dong et al., 2017.

Seeds were surface sterilized using 30% Bleach and sowed on half-strength MS plate (2.2 g/L Murashige and Skoog Basal Salt, 0.5 g/L MES, 8 g/L agar, pH 5.7). The seeds were then stratified at 4 °C for 2 days, induced germination for 3 h under white light, and either put into darkness or 10 µmol/m^2^/s continuous red light for 3-4 days followed by phenotypic observation, RT-qPCR analysis, and western blot analysis.

### Generation of transgenic plants

For generation of *35S: 5U PIF3* and *35S: 5IU PIF3*, *5U PIF3* coding sequence was amplified directly from cDNA, while *5IU PIF3* coding sequence was generated by overlapping PCR of *5IU*, which was amplified from genomic DNA, and *PIF3* coding sequence, which was amplified from cDNA. The resulting *5U PIF3* and *5IU PIF3* coding sequences were inserted into pjim19 (Kan) using XhoI/SacI restriction enzymes. The vectors were then transformed into the *pif3-3* background using *Agrobacterium tumefasciens* strain GV3101 by the floral dip method. Primers used were listed in Supplemental Table 1.

### RT-qPCR

Total RNA was extracted using Qiagen RNeasy Plant Mini kit according to the instructions. Reverse transcription was performed using SuperScript III reverse transcriptase (Invitrogen), oligo(dT) 12-18 (Invitrogen) and 1 µg total RNA as template. The resulting cDNA samples were diluted 10-fold with nuclease-free water, and 2 µl was used for qPCR assay in CFX96 real-time system (Bio-rad). *ACT2* was used as internal control. Primers used for RT-qPCR were listed in Supplemental Table 1.

### Immunoblot

Total protein was extracted from whole seedlings using denaturing buffer (8M urea, 0.1M NaH_2_PO_4_, 0.1M Tris-HCl, pH 8.0). Protein concentration was determined by the Bio-rad protein assay. The same amount of total protein was subjected to SDS-PAGE, transferred to PVDF film, blocked by 5% milk in 1×TBST, and incubated with anti-PIF3 or anti-RPN6 at 4°C overnight. After wash, film was further incubated with HRP-conjugated secondary antibody for 1 h at RT and exposed to X-ray film. The X-ray film was then developed and fixed in a dark room.

### In vitro translation (IVT) assay

For DNA as template, the 5U PIF3 and 5I1U PIF3 coding sequence was inserted into pT7CFE1-Myc (Pierce) vector using NdeI/SacI, yielding pT7-5U PIF3 and pT7-5I1U PIF3, respectively. 1 µg DNA was used as template in a 25 µl in vitro translation system following the instructions of 1-step human coupled IVT kit-DNA (Pierce). The resulting IVT product was mixed with 100 µl loading buffer, boiled, and subjected to western blot analysis using anti-PIF3 antibody. Ponceau S staining was used as loading control. Primers used for construct cloning were listed in Supplemental Table 1.

For RNA as template, the pT7-5U PIF3 and pT7-5I1U PIF3 plasmids were linearized by Not I digestion and purified using QIAquick PCR purification kit. 1 µg linearized plasmid was used as template for in vitro transcription using HiScribe™ T7 High Yield RNA Synthesis Kit (NEB). 5 µg RNA was used as template for IVT, and western blot was done the same as DNA as template.

### Dual luciferase assay

The 35S promoter was cloned into pGreenii 0800-LUC using Hind III/BamH I, yielding pGreenii 35S-LUC. Then the 5’UTR (5U), I_AS_ retained 5’UTR (5IU), uORF mutated 5I_AS_U (5IUm), uORF and mutated uORF (uorf) of PIF3 were cloned into pGreenii 35S-LUC using BamH I/ Spe I. Primers used were listed in Supplemental Table 1.

Protoplasts were isolated based on a protocol previously described (Yoo et al., 2007). The leaves of Col adult plants grown under LD condition for 3-4 weeks were used for protoplast isolation. 10 µg plasmids were used for transfection. After incubated in the dark for 12 h, luciferase activity was measured following the instructions of Dual-Luciferase® Reporter Assay System (Promega). Each plasmid was measured with 4 biological replicates and at least 2 technical repeats.

## Acknowledgements

We thank Prof. Peter Quail and Dr. Weimin Ni for providing *lrb1 lrb2 lrb3* mutant seeds. We also thank Prof. Hongwei Guo for providing *ein3 ebf1 ebf2* mutant seeds. This work was supported by National Institute of Health (GM047850) and National Key R&D Program of China (2017YFA0503800). The authors declare no conflict of interests.

